# New insights into colorectal cancer liver metastasis carcinogenesis and its effect with moxibustion

**DOI:** 10.1101/2021.12.08.471188

**Authors:** Ya-Fang Song, Li-Xia Pei, Jing Guo, Yi Zhuang, Yu-Hang Wang, Ting-Ting Zhao, Lu Chen, Jin-Yong Zhou, Jian-Hua Sun

## Abstract

**Background:** Chemotherapeutic drugs creates severe adverse reactions for colorectal cancer. Moxibustion confers clinical benefits for postoperative patients undergoing chemotherapy, it will fill the blank period of western medicine treatment and provide useful help for tumor patients to prevent recurrence and metastasis, but the physiological mechanisms behind the antitumor effects are unclear. This study was aimed to determine the effect and characterize the differential cytokines and gene expression profiles in intrasplenic transplanted GFP-HCT-116 cells-induced tumors model by Pre-Mox, Post-Mox and Pre-Post-Mox intervention.

**Methods:** Human CRC cells with GFP fluorescence were implanted via intrasplenic injection into Balb/c nude mice spleens. Moxibustion stimulation was applied to the BL18 and ST36 acupoints. The model control (MC) group were given no treatment. Pre-Mox mice were received moxibustion for 2 weeks before HCT-116 cell injection. Post-Mox mice received moxibustion for 3 weeks after CRC cell injection. Pre-Post-Mox mice received moxibustion for 5 weeks (2 weeks before and 3 weeks after CRC cell injection). Peripheral bloods were collected, pooled and assayed using a RayBio® mouse inflammation antibody array. Multi-Analyte Suspension Array was opted for verification. RNA isolated from liver paracancerous tissues from the control group and the experimental groups was subjected to RNA-seq, and then screened out significant differences for in-depth verification.

**RESULTS:** The results showed that moxibustion stimulation increased the survival rate and decreased CRC liver metastasis. With the help of Multi-Analyte Suspension Array and RNA-seq, we screened significant differential expression of cytokines and RNA, then further verified them. The metastasis rate decreased significantly from 100% (10/10, MC group) to 50% (6/12, Pre-Mox group), to 46% (6/13, Post-Mox group), and further to 25% (3/12, Pre-Post-Mox group). Cytokine chips were used significant differences were found in MIP-3α, MDC, IL-6, and IL-1a. Transcriptomic analysis suggested that the low-dose combination of Pre-Mox and Post-Mox modulated larger gene sets than the single treatment. We identified a small subset of genes, like APOA4, IGFBP5, IGFBP6, TIMP1, and MGP, as potential molecular targets involved in the preventive action of the combination of Pre-Mox and Post-Mox.

**Conclusions:** Taken together, the current results provide the first evidence in support of the chemopreventive effect of a combination of Pre-Mox and Post-Mox in CRC. Moreover, the cytokines and transcriptional profile obtained in our study may provide a framework for identifying the mechanisms underlying the carcinogenesis process from colonic cancer to liver metastasis as well as the cancer inhibitory effects and potential molecular targets of Pre-Post-Mox.

## Introduction

Colorectal cancer (CRC) is one of the most common malignant tumors in the world, and is the second highest cause of cancer-related death worldwide^1^. In China, the incidence and mortality of CRC ranked fifth among all malignant tumors, and most of the patients were diagnosed at the middle and advanced stage. Unfortunately, up to 50% of CRC patients either have liver metastases at diagnosis or subsequent metastasis develops soon after, and and this figure has been stable over the last two decades^2, 3, 4^. The age-standardized relative 5-year survival rate of colon cancer patients in China is about 36% - 63%^5^, indicating a long time period from the diagnosis to the final death. This provides an intervention time window for medical workers.

In addition to surgery, chemotherapy and radiotherapy, combined with traditional Chinese medicine is currently the routine choice in the treatment of colon cancer in China. After the completion of chemotherapy for colon cancer patients, in addition to Western medicine regular follow-up, imaging and serological review, combined with the treatment of traditional Chinese medicine, it will fill the blank period of western medicine treatment and provide useful help for tumor patients to prevent recurrence and metastasis. Chemotherapeutic drugs, although toxic to cancer cells, can also induce severe adverse reactions. Moxibustion, an important part of traditional Chinese medicine, has been recognized as a promising treatment for enhancing or coordinating the anti-tumor immune response of the human body in the past 10 years^6, 7, 8^.

Clinical studies have confirmed that moxibustion can ameliorate gastrointestinal disease symptoms, enhance overall body function, and improve the quality of life of postoperative patients undergoing chemotherapy^9^, and also significantly increase the proportion of CD3, CD4, CD8 and NK cells in patients with liver metastasis of colon cancer^10^. While the physiological mechanisms behind the antitumor effects of moxibustion are unclear, our previous research has been suggested that the effect of moxibustion are mediated through innate immune system, leading to reduced the wandering cells from the primary colon cancer via increased influx of neutrophils and macrophages. All these results provide exciting molecular level-evidence indicating the therapeutic effects induced by moxibustion^11^.

Building upon our earlier results, herein we investigate the effects of moxibustion treatment during both the initiation stages of CRC liver metastasis in nude mice. This study aimed to observe the effect of moxibustion on liver metastasis of colon cancer cells, to screen different indexes from serum and liver tissues of nude mice with liver metastasis of colon cancer, and to explore its possible mechanism from the aspects of inhibition of metastasis and anti-tumor respectively.

## Materials and Methods

### Study design

To investigate the effects and mechanisms of moxibustion in colon cancer liver metastasis, BALB/C mice were randomly divided into four groups: G1: Model control group (MC, n = 15); G2: Pre-Mox group (2 weeks of pretreatment with moxibustion before CRC cell injection, n = 15); G3: Post-Mox group (3 weeks of moxibustion treatment after CRC cell injection, n = 15); G4: Pre-Post-Mox group (2 weeks of moxibustion treatment before and 3 weeks of moxibustion treatment after, CRC cell injection, n = 15). Infected HCT 116 cells with GFP fluorescence were implanted via intrasplenic injection to establish the murine orthotopic CRC model. Moxibustion treatment at bilateral Ganshu (BL18) and bilateral Zusanli (ST36) acupoints were applied on mice at different period times in three groups (once daily for 10 min, seven times a week). Model control mice were fixed at the same time without any treatment. The experimental workflow is shown in Figure 1.

**Figure 1.**
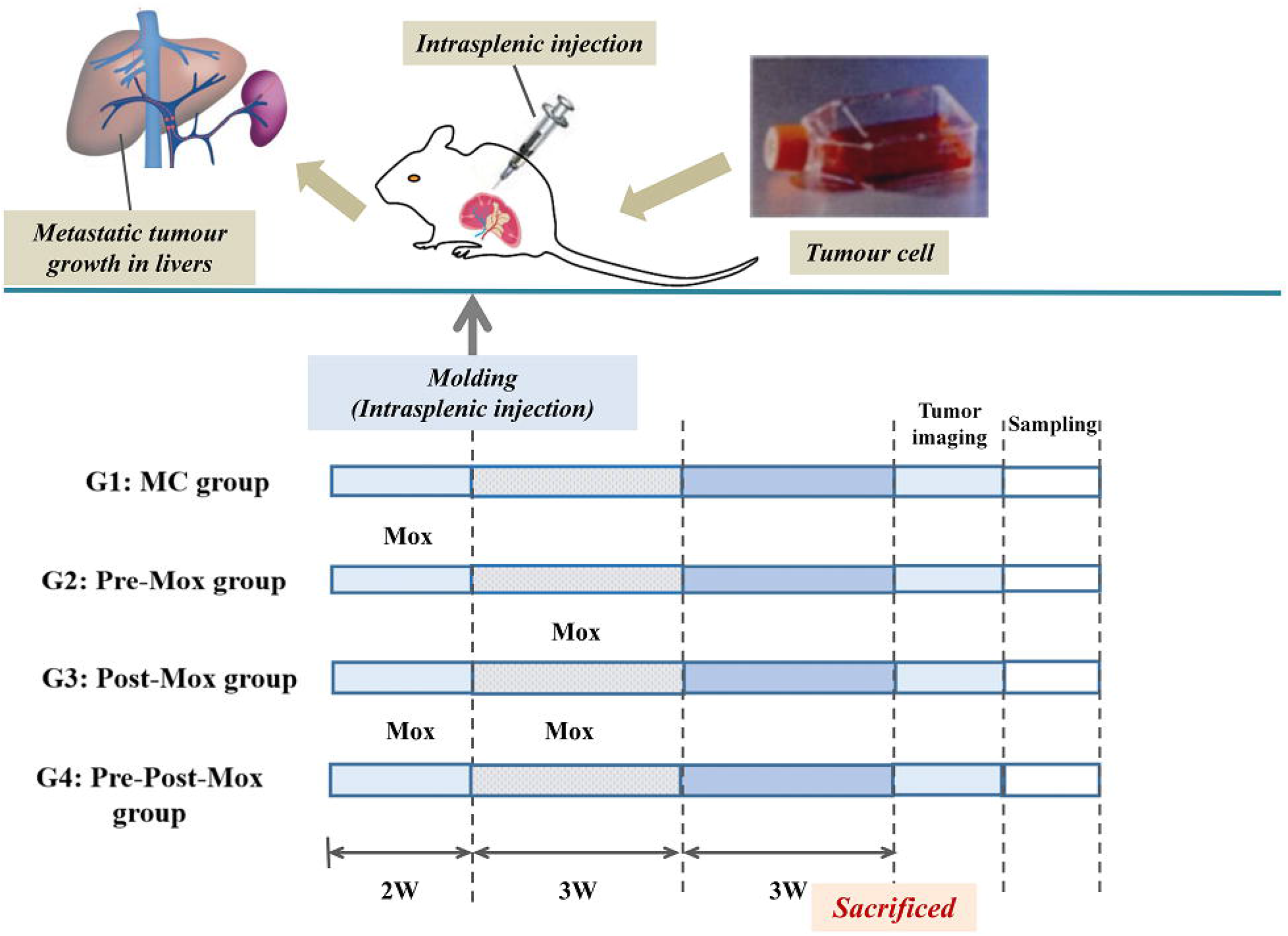
The experimental workflow: Tumor formation and moxibustion intervention timeline for G1, G2, G3 and G4 group were as follows. All nude mice were administered with 20 μL cell suspension in the spleen. G1: MC group, G2: Pre-Mox group, G3: Post-Mox and G4: Pre-Post-Mox group.

### Animals

A total of sixty male BALB/C mice (20∼24g, aged 4∼6 weeks) were purchased from Comparative Medicine Centre of Yangzhou University (Yangzhou, Jiangsu, China). All mice were housed at an ambient temperature of 24°C with a 12 h light-dark cycle in the SPF animal houses. The feed adaptation lasted for seven days. Research was conducted in accordance with institutional guidelines and approved by Animal Ethics Committee of Affiliated Hospital of Nanjing University of Chinese Medicine (Ethical review code: 2018 DW-03-03).

### Model establishment: Intrasplenic injection

GFP-HCT-116 cells were cultured in1640 with 37 °C incubator and 5% CO2, and collected to the tumor cells concentration of 1×10^8^/mL. Giavazzi R’s method was used to intrasplenically transplant cancer cells12. Briefly, all mice were anesthetized with 25% chloramine ketone(1ul/g, i.p.). GFP-HCT-116 cells were implanted via intrasplenic injection into sixty nude mice (2 × 10^6^ cells/mouse in 20 μ l of physiological solution). 1 cm laparotomy was performed in the left subcostal region of the abdomen. The spleen was gently exposed and the cells were injected. The spleen was then put back into the abdominal cavity and the abdominal wall was sutured with 5.0 stitches. They were recovered on a heating pad after surgery, till being returned to housing cages. The liver and spleen were removed and cleaned, followed by careful examination of the tumors. After removing the proximal end and palpable tumors, the remaining normal portion of the liver and spleen were snap frozen and stored at -80 ° C for molecular assays. At the end of the experiment, normal spleen tissue, spleen primary tumor, normal liver tissue and liver primary tumor were taken and weighed, and the tumor formation rate and liver metastasis rate among the groups were calculated. The number of tumors, tumor weight and the number of metastatic foci could be seen. Macroscopically, the degree of primary tumor formation and liver metastasis among each group was evaluated. The status, food in-take and weight change of nude mice were observed twice one week. At the end of the experiment, the mice were killed, and the liver and spleen were taken out to investigate the degree of liver metastasis. We evaluated metastatic liver tumors macroscopically by counting visible tumor number and measuring m aximal size of tumors, area. Tumor images were acquired by a fluorescence stereo-microscope.

### Treatment: Moxibustion

The mice in the G2, G3 and G4 groups were treated with moxibustion at bilateral BL18 and ST 36 acupoints. The locations for these acupoints were determined according to Government Channel and Points Standard GB12346-90 of China and “ The Veterinary Acupuncture of China.” The moxa stick (diameter 4mm, length 12 cm) was ignited and placed 1.5 cm above the selected points for 10 min per day. The whole treatment course varied at different period times in three groups. The G1 group was given the same fixation (the tail of the mice were fixed on the fixation device) as the G2, G3 and G4 group without any intervention.

### Animal plasma collection for RayBiotech mice cytokines arrays

Peripheral blood was collected from each mouse in the four groups at the time of sacrifice. Plasmas within the same group were pooled and assayed using a RayBio® mouse inflammation antibody array(Raybiotech, QAM-CYT-5, Norcross GA, USA) according to the manufacturer’s instructions^13^. Each representing a unique set of 40 antigen-specific antibodies to detect a total of 200 markers on a glass slide matrix (Additional file 1). To increase the accuracy of the measurement, three replicates per antibody were spotted, and the averages of the median signal intensities across replicate spots (minus local background) were used for all calculations. After development, films were scanned and the images processed and quantified using the Normalization data. The differential proteins were screened by fold change. The selection conditions were as follows: 1) fold change = < 0.83 or fold change > = 1.2. 2) The average (fluorescence) signal value of each group was > 150.

### Multi-Analyte Suspension Array (MASA)

Subsequently, biotinylated antibody labeling was performed using MCYTOMAG-70K-06 and MECY2MAG-73K-03 kit, according to the manufacturer’s instructions. The detection volume of each sample is not less than 100ul. As is subsequently described, MASA was used to examine the IL-1α, KC, LIX, MCP-1, IL-6, TNF-α, MDC, MIP-3α and MIP-3β expression in G2, G3 and G4 groups, and G1 mice without any intervention.

### RNA sequencing

RNA was purified from liver paracancerous tissues collected from the intrasplenic injection-induced tumors in BALB/C mice. RNA-Seq was performed using Illumina HiseqTM2000 sequencer. The original RNA SEQ data were output in fastq format. The direct expression of gene expression level is the abundance of transcripts, which is quantified by stringtie. A cut-off value of a log2 fold change greater than 2 and an adjusted p value (q value) less than 0.05 were used to extract the differentially expressed genes.

### Quantitative polymerase chain reaction (qPCR)

The paracancerous tissues of liver metastases were extracted by real-time fluorescence quantitative PCR (FQ-PCR). After total RNA extraction, RNA agarose gel electrophoresis was performed, and reverse transcripts of cDNA, were selected according to the requirements of Real-Time PCR primers, followed by cDNA and primer quality detection. The expression of target gene was analyzed, and the amplification curve and dissolution curve were formed by quantitative PCR.

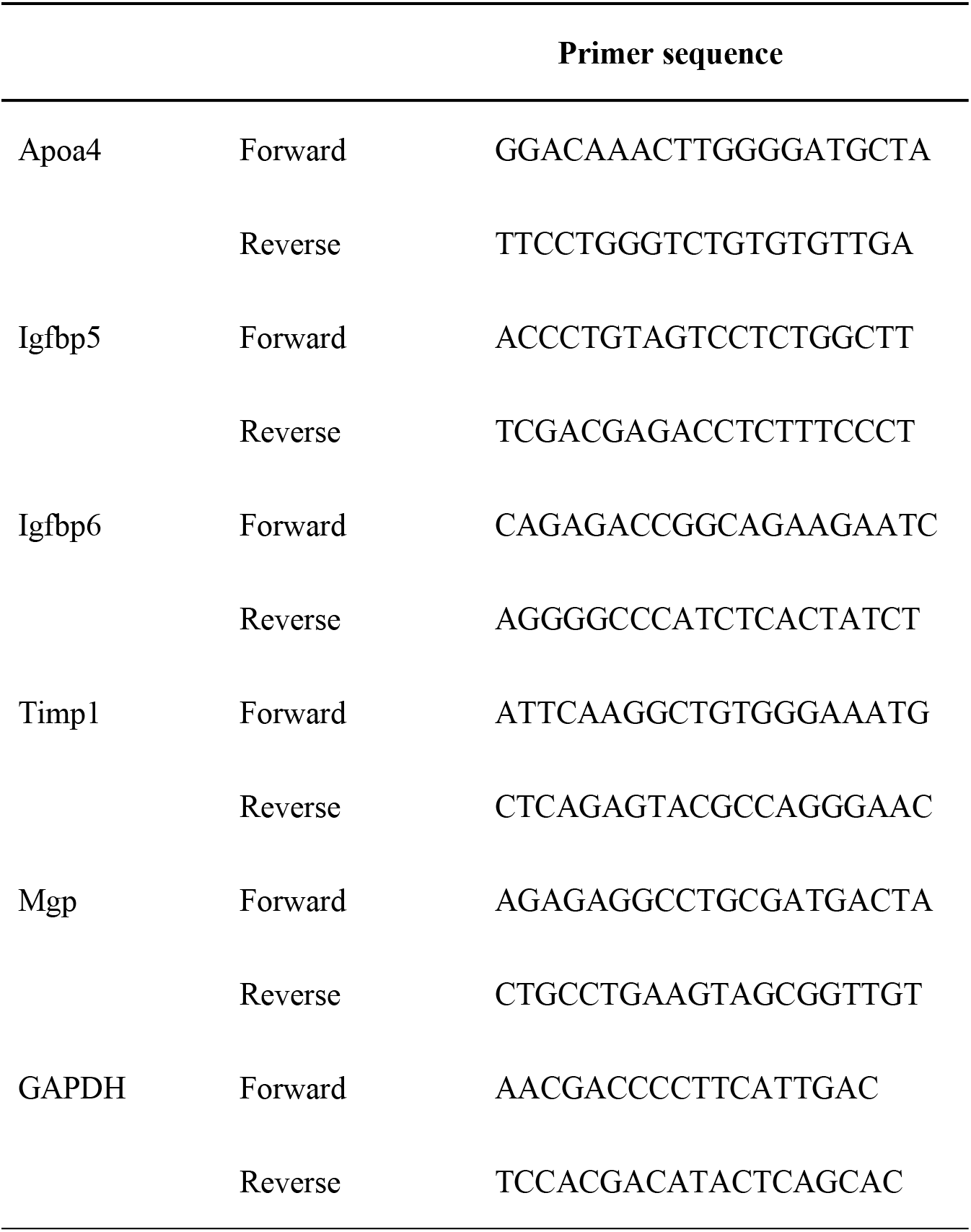

### Statistical analyses

Statistical analyses were performed using SPSS 24.0 software (SPSS Inc, Chicago, Illinois, USA). Data were expressed as mean ± SD (Standard ± Deviation). One-way analysis of variance (ANOVA) was performed for normally distributed data and a non-parametric test (Kruskal–Wallis H test.) was used for non-normally distributed data. Differences in relative abundance between groups were assessed using the Kruskal–Wallis test. *P < 0*.*05* was considered statistically significant.

## Results

### Effects of Pre-Mox, Post-Mox and their combination in the prevention of colon cancer liver metastasis

After the mice were sacrificed, the situation of colon cancer liver metastasis was examined by a fluorescence optical imaging system. As shown in Figure 2, green clusters represent tumor tissue. Strict criteria were used to determine whether liver metastasis occurred in mice, that is, there was no green dot in the captured liver image. When taking moxibustion, the metastasis rate of the MC group was 100%, indicating a successful model; the liver metastasis rate decreased from 100% (10/10, MC group) to 50% (6/12, *P* = 0.009, Pre-Mox group), 46% (6/13, *P* = 0.005, Post-Mox group), and then reduced to 25% (3/12, *P* = 0.000, Pre-Post-Mox group), the latter three groups were significantly different from the MC group(Figure3(a)). Compared with the MC group, the number of liver metastases tumor, spleen tumor volume, and spleen tumor weight in the Pre-Mox group, Post-Mox group and Pre-Post-Mox group displayed significantly decreased expression (P < 0.01), though the differences are not statistically significant among the Pre-Mox group, Post-Mox group and Pre-Post-Mox group (P > 0.05) (Figure3(b)-(d)). To the liver tumor weight, a similar trend was observed between the MC group and the Pre-Post-Mox group (P < 0.001, Figure3(e)). These findings suggest that moxibustion may have a better effect in inhibiting liver metastasis of tumor cells, and the Pre-Post-Mox group has the lowest rate of liver metastasis in nude mice.

**Figure 2:**
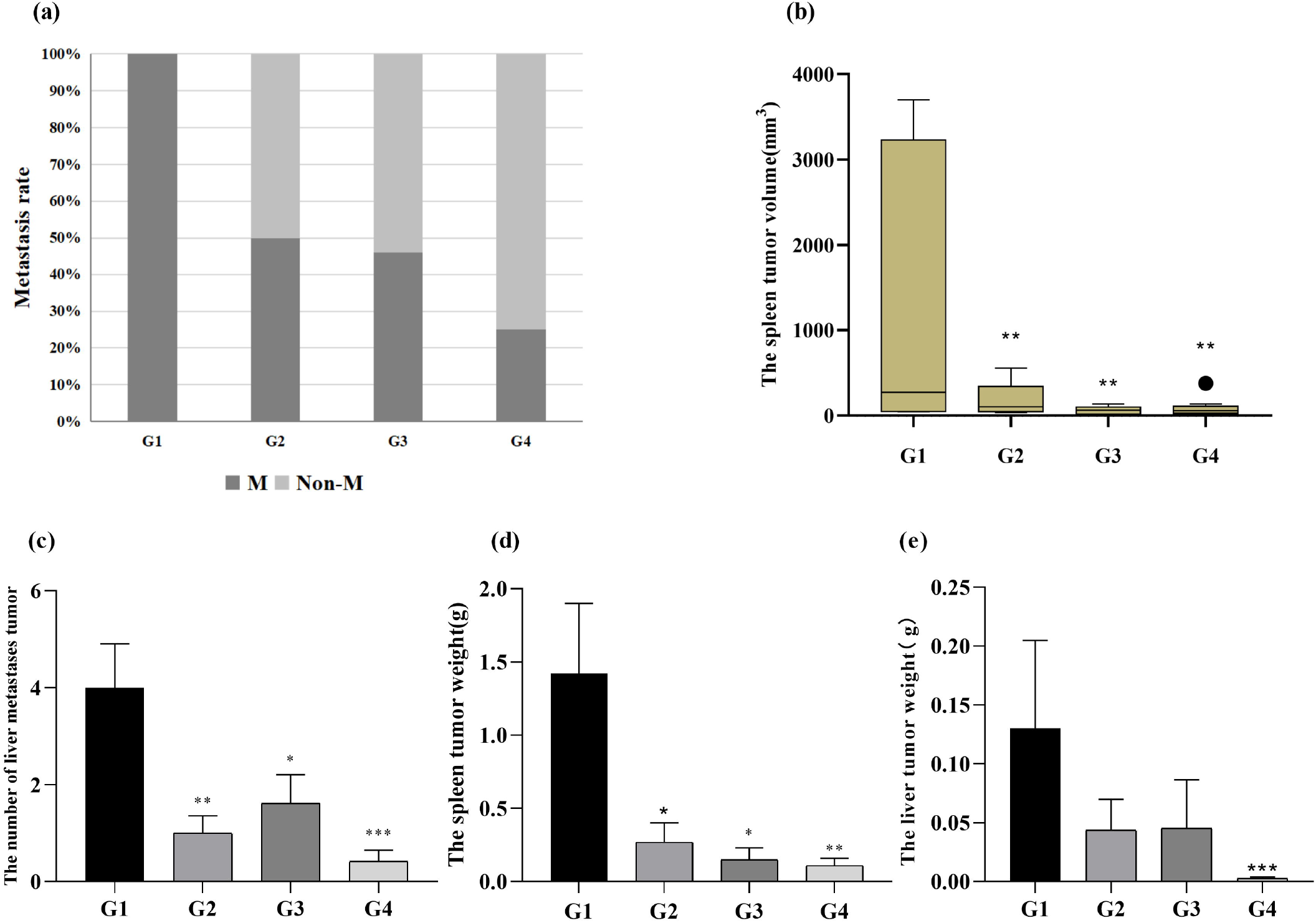
Representative images of CRC liver metastasis. A–D: Images of mice anatomy showing the inner organs. The white left arrows indicate the transplanted tumor in the spleen. E-H: Liver Images. The down arrows indicate the metastatic tumors. A, E: G1 - MC group; B, F: G2 - Pre-Mox group; C, G: G3 - Post-Mox group; D, H: G4 - Pre-Post-Mox group.

**Figure 3:**
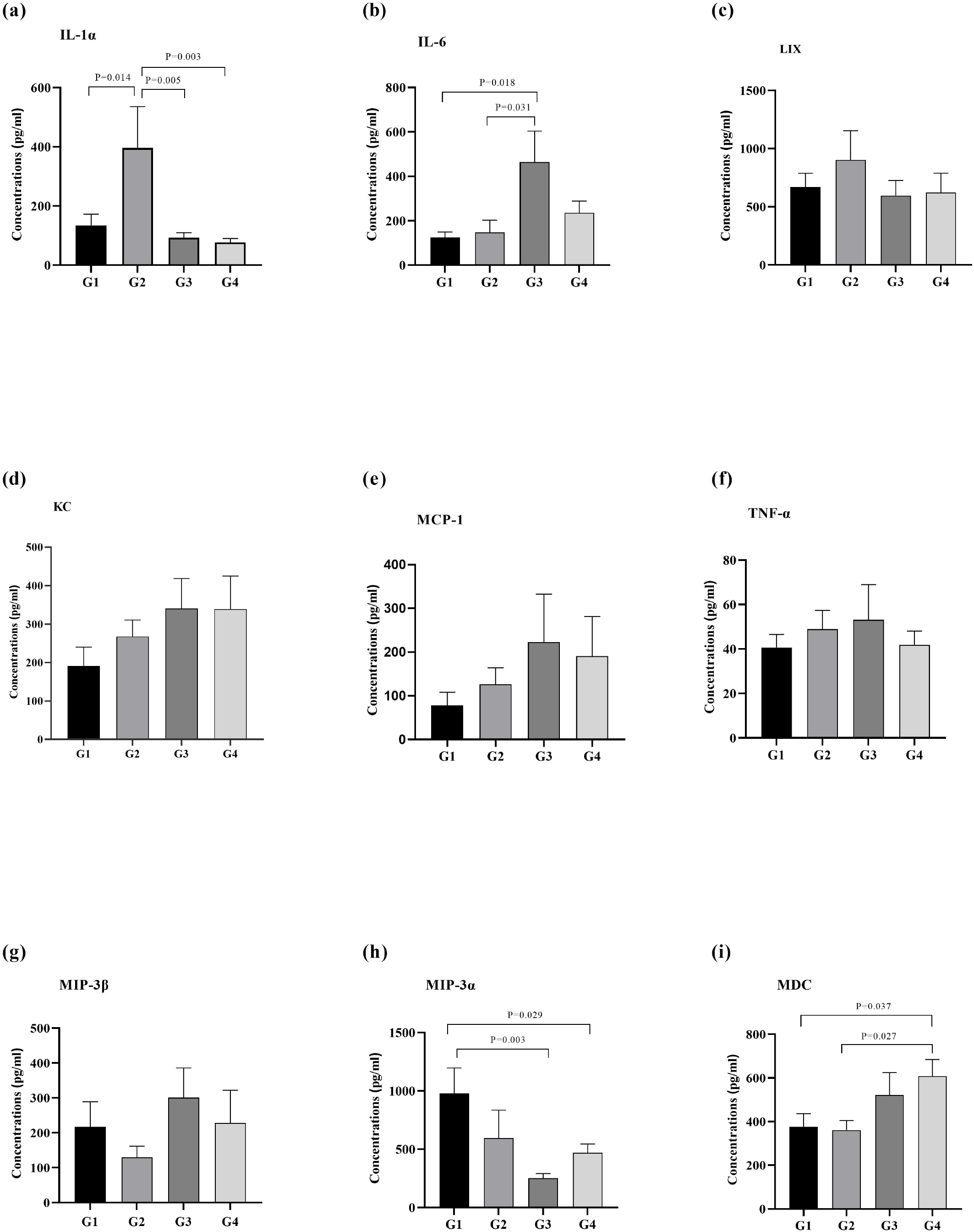
Moxibustion’s inhibitory effect on CRC liver metastasis.*P < 0.05; **P < 0.01, ***P < 0.001.

### Identification of Potential Targets expression in the peripheral blood of nude mice

Among the 200 cytokines detected by GSM-CAA-4000 antibody chip, compared with the MC group, 77 kinds of cytokines in the serum of nude mice with liver metastasis of colon cancer cells in the Pre-Mox group met the standard of differentially expressed protein <u>(supplement 1)</u>, 93 kinds of cytokines in the Post-Mox group met the standard of differentially expressed protein <u>(supplement 2)</u>, and 94 kinds of cytokines in the Pre-Post-Mox group met the standard of differentially expressed protein <u>(supplement 3)</u>. To further explore the role of these differentially expressed cytokines in moxibustion on inhibiting liver metastasis of colon cancer cells, we screened chemokines IL-1α,KC,LIX, MCP-1,IL-6,TNF-α,MDC,MIP-3α,MIP-3β, and then verified them by liquid phase microarray (Table 1).

**Table 1:**
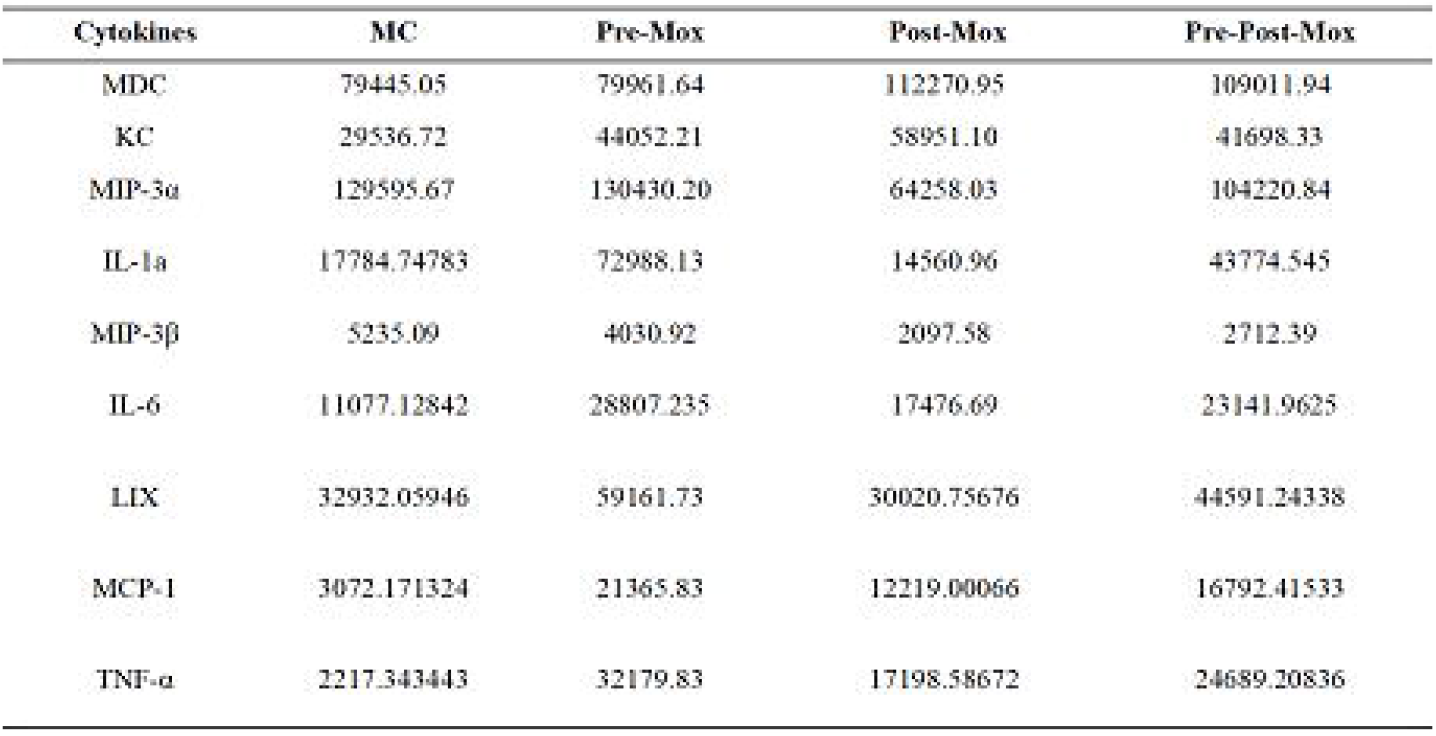
Changes of cytokines in mice plasma after moxibustion treatment (fluorescence signal value.

IL-1α, IL-6, MDC and TNF-α are main cytokines secreted by macrophages. We next evaluated whether these are affected by moxibustion. As shown in Figures 4(a), compared with the MC group, the IL-1α expression levels the Pre-Mox group displayed significantly increased expression (P < 0.01). A similar trend was observed in the Post-Mox group with the IL-6 expression levels being higher (Figures 4(b)). LIX, KC, MCP-1 and TNF-α play a key role in the regulation of tumor microenvironment and metastasis, and are closely related to the evolution, implantation, proliferation and inflammatory cell infiltration of colon cancer. However, LIX, KC, MCP-1 and TNF-α, were observed no significantly difference after moxibustion intervention(P > 0.05, Figures 4(c)-(f)). MIP-3α and MIP-3β are important downstream molecules for immune activation. Compared with the MC group, the expression level of MIP-3 α in Post-Mox group and Pre-Post-Mox group decreased significantly (P < 0.01), a similar trend was observed in the Pre-Mox group without statistically difference (P > 0.05, Figures 4(h)). However, we didn’ t observe significantly difference with the MIP-3β expression levels after moxibustion intervention (P > 0.05, Figures 4(g)). In our study, we also found that the level of MDC increased significantly in the Pre-Post-Mox group compared with the MC group.A similar trend is observed in the Pre-Mox group and the Pre-Post-Mox group, with the MDC expression levels being lower in the Pre-Mox group than in the Pre-Post-Mox group (P < 0.05, Figures 4(i)).

**Figure 4:**
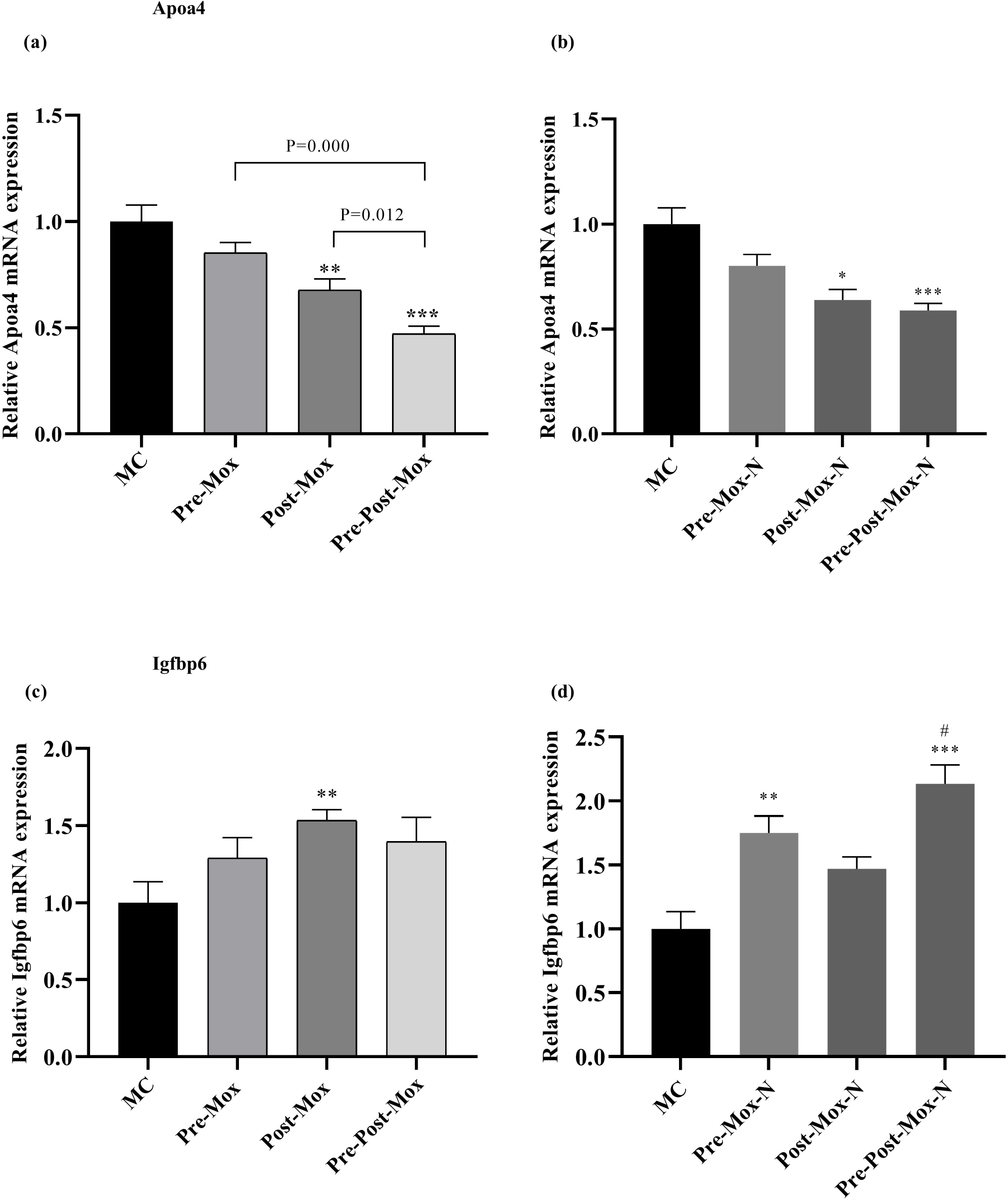
Analysis of cytokine concentration expression level. (a): IL-1α expression levels; (b): IL-6; (c): LIX expression levels; (d) KC expression levels; (e) MCP-1 expression levels; (f): TNF-α expression levels; (g): MIP-3β expression levels; (h): MIP-3αexpression levels; (i): MDC expression levels.

### Overview of differentially expressed genes regulated by the treatment of Pre-Mox, Post-Mox and their combination compared to the MC group

To determine how Pre-Mox, Post-Mox and their combination exerted a preventive effect in the intrasplenic transplanted GFP-HCT-116 cells-induced colon cancer liver metastasis model, we compared the global gene expression profiles of paracancerous tissue of liver tumors to those treated by Pre-Mox, Post-Mox, or their combination Pre-Post-Mox. As shown in Figure5(a), we observed 17 differentially expressed genes in comparison with liver tumor-adjacent tissue from Pre-Mox treated mice versus tumors and the model group (7 genes were up-regulated, while 10 genes were down-regulated), we also observed 51 significant genes in the Pre-Mox mice without liver metastasis liver metastasis compared with the MC group (27 up-regulated genes and 24 down-regulated genes). In the Post-Mox treatment, we observed 42 genes with differential expression in comparison with the model group (10 genes were up-regulated by Post-Mox treatment, while 32 genes were down-regulated). We conducted an in-depth study and identified 107 genes with differential expression in comparison with paracancerous tissues from Post-Mox treated mice with no liver metastasis and the model group (16 genes were up-regulated by Post-Mox treatment, while 91 genes were down-regulated). The combination of Pre-Post-Mox treated was found to modulate more genes than Pre-Mox or Post-Mox alone when compared to the MC group. A total of 47 genes that showed significantly differential expression levels were identified (23 genes were up regulated while 24 genes were down-regulated). We also observed 106 genes with differential expression in comparison with liver tumor-adjacent tissue from Pre-Post-Mox treated mice without liver metastasis versus tumors and the MC group (31 genes were up-regulated by Post-Mox treatment, while 75 genes were down-regulated). This result indicated that 3 weeks of moxibustion treatment after CRC cell injection (Post-Mox treatment) modulated a larger number of genes than 2 weeks of pretreatment with moxibustion before CRC cell injection (Pre-Mox treatment), whereas the combined treatment (Pre-Post-Mox treatment) was able to regulate an even broader gene set.

**Figure 5:**
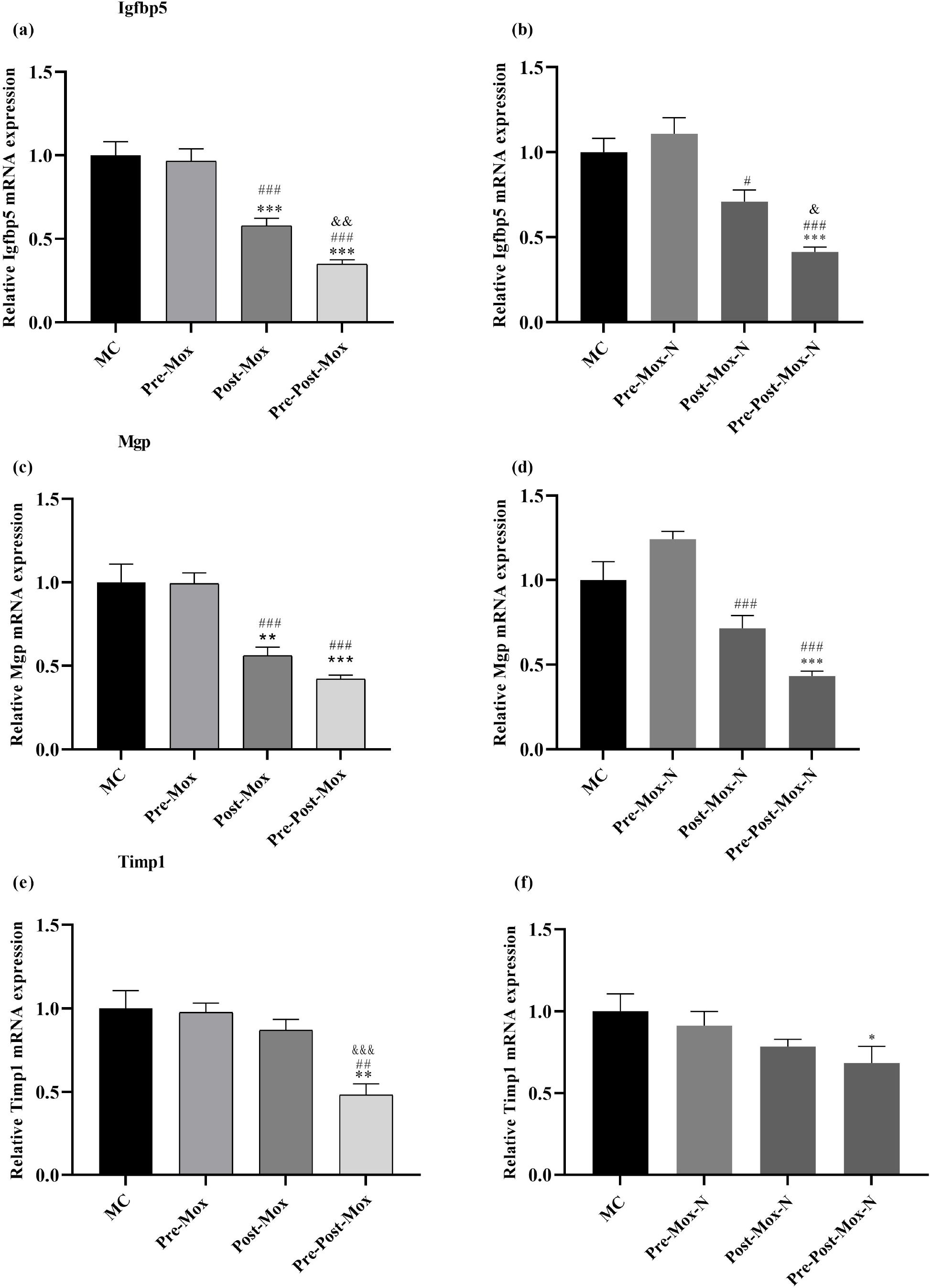
Overview of differentially expressed genes regulated in the four group.(a) Statistics analysis of gene expression differences among four groups; (b) Principal coordinate analysis (PCoA) among the four groups (weighted analysis); (c): differential genes venn diagrams; (d):Co-occurrence matrix of clustering mice. The more similar the mycobiome, the higher the tendency to cluster together.

In our study, we identified a total of 5 genes that appeared in all three treatment groups (Pre-Mox, Post-Mox, and their combination) compared to the MC group. In addition, we observed a total of 8 genes that appeared in all three treatment mice (Pre-Mox, Post-Mox, and their combination) without liver metastasis compared to the MC group (Figure5(b)). Principal coordinates analysis (PCoA) demonstrated significant differences among Pre-Mox, Post-Mox, and their combination compared to the MC group on the first axis (Figure5(c)). However, the first and second principal coordinates accounted for 56.25% and 24.8% of th e total variations respectively, indicating that moxibustion may inhibit liver metastasis of colon cancer. The grayer color is represented the higher the similarity. The yellower color is represented the the lower the similarity. As shown in the heat map (Figure5(d)), the color of the combination group was overall slightly yellow color than that of the Pre-Mox and Post-Mox groups, suggesting that combined treatment with Pre-Mox and Post-Mox groupsmay regulate the expression of these shared genes at a higher fold change than single treatment.

### Top differentially expressed genes by Pre-Mox, Post-Mox and their combination compared to the MC group

Upon further examination of the significant differentially expressed gene profiles of liver tumor-adjacent tissue, we screened out Apoa4, Igfbp6 genes that appeared significant differences in at least two of all three treatment groups (Pre-Mox and Post-Mox, and their combination) compared to those in the MC group (Table 2). As shown in Figure 6(a)-(b), the expression of Apoa4, Igfbp6 was validated in liver tumor-adjacent tissue by qPCR analysis, confirming that its expression was altered by these treatments and that the combination was able to decrease its expression to a higher magnitude. A similar trend was observed in the three treatment mice (Pre-Mox, Post-Mox, and their combination) without liver metastasis compared to the MC group. Similarly, Igfbp5 and Mgp were the top 2 down-regulated genes in all three treatment mice (Pre-Mox, Post-Mox, and their combination) without liver metastasis compared to the MC group (Table 2), and this trend was also confirmed by qPCR analysis (Figure 7(a)-(d)). In addition, among the top 10 up or down-regulated genes in the Pre-Post-Mox treated group without liver metastasis, the expression of Timp1 genes was more than 100-fold lower than that in the liver tumor-adjacent tissue from the Pre-Mox/Post-Mox group, and this trend was also confirmed by qPCR analysis (Figure 7(e)-(f)). These results show that the combination Pre-Post-Mox treatment effectively prevented liver metastasis carcinogenesis in our study.

**Table 2:** The q values of significant differences genes. qValue: p value after correction. The smaller the q value, the more significant difference in gene expression. N: non-metastasis.

| Gene Name | qValue |  |  |  |  |  |
| --- | --- | --- | --- | --- | --- | --- |
|  | G2 vs G1 | G2-N vs G1 | G3 vs G1 | G3-N vs G1 | G4 vs G1 | G4-N vs G1 |
| Apoa4 | 0.0425228 | 2.08E-06 | 0.000788225 | 0.002144714 | 0.010979361 | 0.0425228 |
| Igfbp5 | 0.00027924 | 0.000257254 | 0.000660458 | 4.22E-09 | 1.34E-06 | 7.35E-18 |
| Igfbp6 | 0.014929186 | 0.020744939 | 1.56E-05 | 0.000201439 | 0.017453038 | 2.28E-07 |
| Mgp | 0.001630289 | 0.006841453 | 2.24E-21 | 5.13E-09 | 0.017453038 | 4.81E-06 |
| Timp1 | 0.025159337 | 0.000110087 | 0.016600418 | 0.002670022 | 0.015304551 | 6.78E-07 |

**Figure 6:**
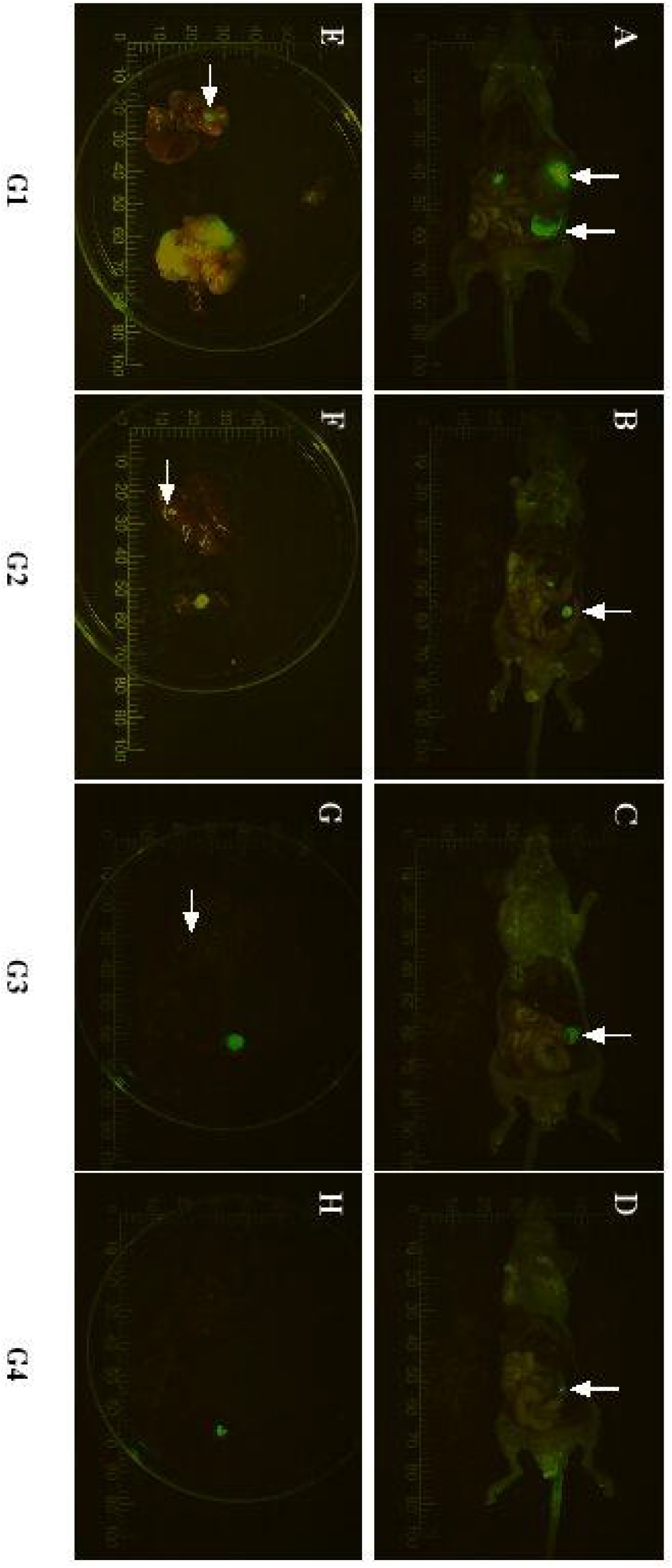
mRNA expression levels in the liver adjacent tissues. (a), (b) Apoa4 mRNA levels; (c),(d): Igfbp6 mRNA levels. N: non-metastasis. *P < 0.05; **P < 0.01, ***P < 0.001.

**Figure 7:**
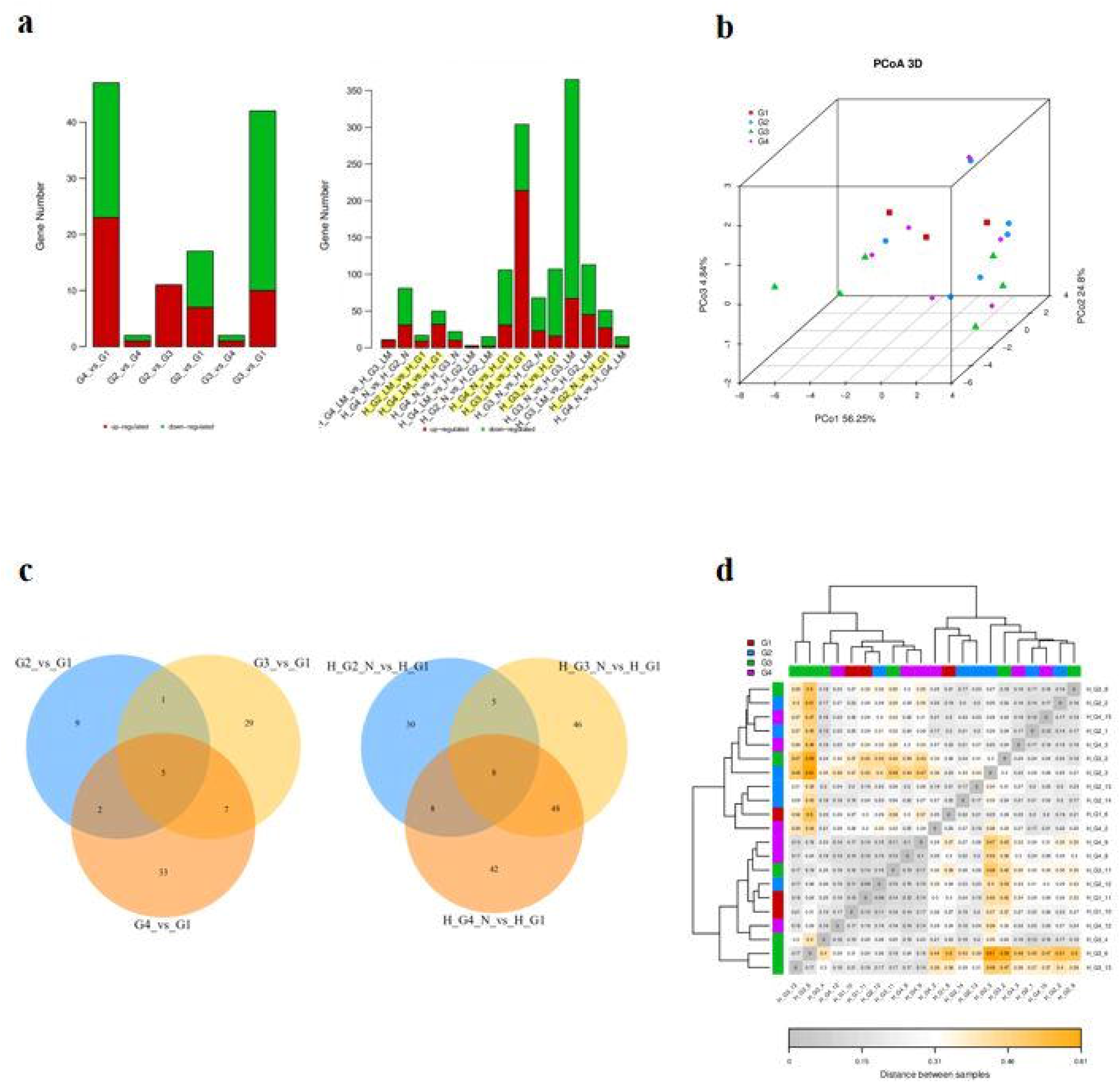
mRNA expression levels in the liver adjacent tissues. (a), (b) Igfbp6 mRNA levels; (c),(d): Mgp mRNA levels. (e), (f) Timp1 mRNA levels. *P < 0.05; **P < 0.01, ***P < 0.001.

## Discussion

On the whole, the pitching-in opportunity of traditional Chinese medicine in the treatment of CRC in China is still relatively late, and few patients take in the early stage. The results of the present study showed that moxibustion stimulation could inhibit the liver metastasis rate, reduce the mortality, improve the microenvironment and Pre-Post-Mox treatment had the best effect.

Inflammation is an important feature of tumors^14^. Cytokines, as a catalyst to other effector cells, are an important type of biological mediator, which are secreted by a variety of immune cells and exist in the peripheral blood. With the goal of uncovering biomarkers indicative of liver metastasis, we opted for a large multiplex antibody array that could simultaneously probe blood samples for 200 proteins and finally screened out significant differences IL-1α, IL-6, LIX, KC, MCP-1, TNF-α, MIP-3β, MIP-3α, and MDC for further verification. When evaluating these nine markers, we noted that they were not all associated with one common process or pathway, but instead spanned across multiple pathways in the huge network regulating tumor occurrence, metastasis, invasion, angiogenesis. Liquid chip technology was used to detect nine cytokines in the peripheral blood plasma for further verifing and the results showed that the levels of MIP-3α, MDC, IL-6, IL-1a significantly higher than those of model control nude mice. MIP-3α, as a new target for colon cancer liver metastasis and other metastatic processes, has potential clinical value for the staging of primary tumors and the prevention of liver metastasis recurrence^15^.The high expression of MIP-3α in liver metastases is related to the malignant degree of metastatic cells^16, 17, 18^. MDC (also known as CCL22) is mainly secreted by macrophages, natural killer (NK) cells, T cells, dendritic cells. CCL22 can control T cell immunity by recruiting T-regs from tumor tissue and promoting the formation of DC-T-reg contact in lymph nodes^19^. However, the T-cell-deficient Balb/c nude mice were used in this study. In this case, the role of macrophages and NK cells were more important. It is somewhat similar to the first line of defense of the body in tumor metastasis. The metastatic cells were eliminated in the budding state and reduced the formation of the final metastasis, and the specific mechanism needs to be further investigated. IL-1α and pro-inflammatory IL-6 cytokines secreted by macrophages are closely related to colon cancer cell proliferation, apoptosis and metaplasia^20, 21^, which can mediate immune and inflammatory reactions and inhibit cell apoptosis in an inflammatory environment. In addition, LIX, MIP-3β, MCP-1, TNF-α, and KC also showed an increasing trend, which may also play an anti-tumor effect. Such multiple cytokine biomarkers, while of obvious interest from a intervention effect perspective, could also provide some interesting insight into the cause and development of CRC.

With the help of RNA-seq, we established, for the first time, a global transcriptome profile associated with Pre-Mox, Post-Mox, and their combination Pre-Post-Mox. Finally, we screened APOA4, IGFBP5, IGFBP6, TIMP1, and MGP genes with significant differences, and then conducted in-depth verification. Previous studies on Apolipoprotein A-IV (Apoa4) were mainly focused on anti-inflammatory^22^, antioxidant^23^, lipid metabolism^24^, etc. At present, the research on the involvement of APOA4 in cancer mainly focused on ovarian cancer. Studies have showed that Apoa4 is in serum of ovarian cancer. However, since there is no evidence that ApoA-IV is expressed in large amounts in ovarian tissue, it is not clear why ovarian cancer causes a decrease in circulating ApoA-IV levels^25^. Insulin-like growth factor-binding protein-5 (IGFBP-5) are considered to be critical players in colon cancer-induced liver metastasis^26, 27, 28^. Overexpression of IGFBP-5 indicates poor prognosis. Previous studies have shown that it has inhibitory effect in colon cancer^29^, breast cancer^30^, urothelial carcinoma^31^, neuroblastoma^32^ and osteosarcoma^33^. Tissue inhibitor of metalloproteases1(TIMP1) expression is significantly up-regulated in colon cancer, positively correlated with the size, lymphatic metastasis and clinical stage while negatively correlated with the survival time of patients^34^. Matrix Gla protein (MGP), a marker of colorectal cancer, is a vitamin K-dependent extracellular matrix protein^35^.

The occurrence of CRC liver metastasis is a multi-step and selective process, and its regulation mechanism is very complex. The synergistic effect of different genes and different signal pathways regulates the key steps of CRC liver metastasis. The transcriptional level suggests thought and theoretical basis for exploring the mechanism of moxibustion inhibiting liver metastasis. Here, we report that moxibustion intervention, especially concomitant administration of 2 weeks of moxibustion treatment before and 3 weeks of moxibustion treatment after CRC cell injection, effectively attenuates tumor growth in the liver and improve the microenvironment by down-regulating the transcriptional levels of Apoa4, Igfbp5, Timp1 and Mgp, and up-regulating Igfbp6.

However, there were some limitations in this study. Firstly, since the weight of liver metastases in this experiment were too light (< 0.01g), not enough at least 3 samples in each group. In order to reduce the error and difference, only the adjacent tissues were taken. Secondly, our experiment explored for a long time in the early stage, including the selection of animal models, intervention methods, acupoints and intervention weeks. The current results show a good start, but only screened out the genes or proteins with significantly differences from the gene and protein levels. It has no in-depth exploration on these genes or proteins. Thirdly, further studies comparing the preventive effect of moxibustion against liver metastasis in different strains of animals and different ethnicities of patients should be considered.

## Conclusions

In summary, our study is the first to show that moxibustion administration effectively attenuates tumor metastasis. Our results provide a quantitative cytokines and proteins expression profile of CRC cell -induced tumors as well as tumors from nude mice treated with Pre-Mox, Post-Mox, and their combination. Furthermore, a small set of cytokines and proteins were postulated as potential molecular targets involved in the action of Pre-Post-Mox treatment in the prevention of CRC cell-induced model. These findings provide novel insights that further the understanding of the carcinogenesis of CRC liver metastasis as well as the mechanisms underlying the preventive effect of moxibustion treatment.

## Author Contributions

We thank Dr. Yi Zhuang for her assistance in the histology evaluation. We also thank Dr. Jinyong Zhou for their help in the animal study. The authors are grateful to Nanjing Yuanduan Biotechnology Co., Ltd for providing GFP-HCT-116 cells. We thank all of the members of Dr. Jianhua Sun’ s team for their helpful discussions and the preparation of this manuscript. We thank the management of Dongnan University for providing the required infrastructure to complete the research work.

## Funding

This work was supported by 81473748 and 81804193 from the National Natural Science Fund, y2018rc05 and y2018rc25 from the Peak Talent Project of Jiangsu Province Hospital of Chinese Medicine.

## Competing Interests

All authors declare there are no competing interests

## Ethics Statement

The experimental protocol was approved by the Animal Ethics Committee of the Affiliated Hospital of Nanjing University of Chinese Medicine and implemented strictly according to the experimental protocol (Ethical review code: 2018 DW-03-03).

### Patient consent for publication

Not required.

### Provenance and peer review

Not commissioned; externally peer reviewed.

## Data Availability Statement

The raw data supporting the conclusions of this article will be made available by the authors, without undue reservation. Further inquiries can be directed to the corresponding authors.

## Supplemental material

This content has been supplied by the author(s). It has not been vetted by BMJ Publishing Group Limited (BMJ) and may not have been peer-reviewed. Any opinions or recommendations discussed are solely those of the author(s) and are not endorsed by BMJ. BMJ disclaims all liability and responsibility arising from any reliance placed on the content. Where the content includes any translated material, BMJ does not warrant the accuracy and reliability of the translations (including but not limited to local regulations, clinical guidelines, terminology, drug names and drug dosages), and is not responsible for any error and/or omissions arising from translation and adaptation or otherwise.

## Open access

This is an open access article distributed in accordance with the Creative Commons Attribution Non Commercial (CC BY-NC 4.0) license, which permits others to distribute, remix, adapt, build upon this work non-commercially, and license their derivative works on different terms, provided the original work is properly cited, appropriate credit is given, any changes made indicated, and the use is non-commercial. See http://creativecommons.org/licenses/by-nc/4.0/.

## Notes

### Competing Interest Statement

The authors have declared no competing interest.

