## Supplementary figures and images for "New insights into colorectal cancer liver metastasis carcinogenesis and its effect with moxibustion"

### file1

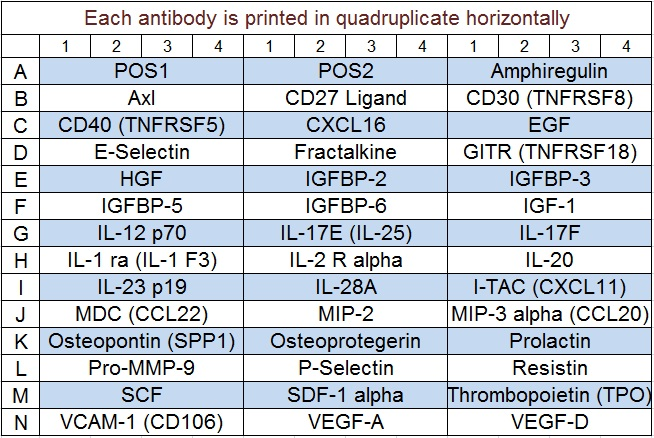

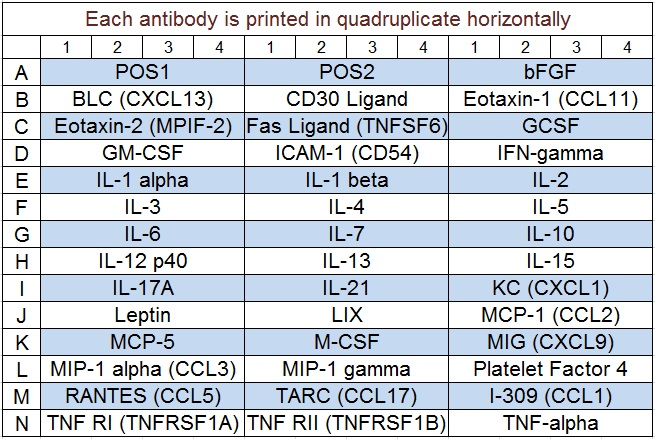

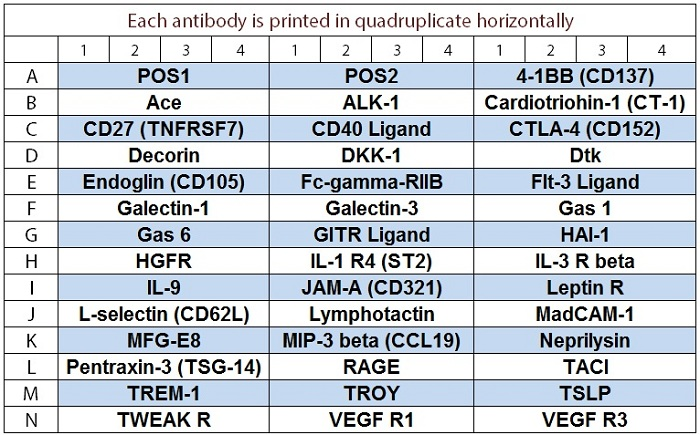

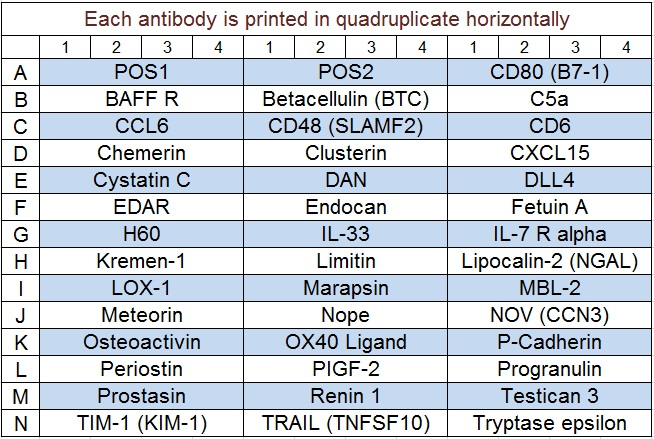

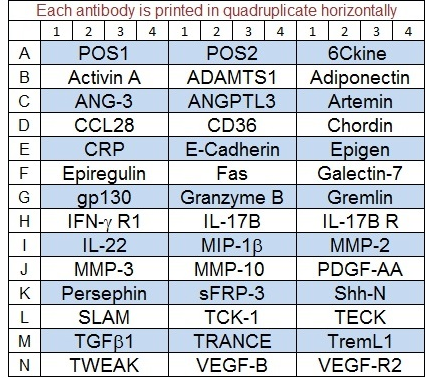
